# Microbubble dynamics in brain microvessels

**DOI:** 10.1101/2023.10.25.563941

**Authors:** James H. Bezer, Paul Prentice, William Lim Kee Chang, Sophie V. Morse, Kirsten Christensen-Jeffries, Christopher J. Rowlands, Andriy S. Kozlov, James J. Choi

## Abstract

**Background:** Focused ultrasound stimulation of microbubbles is being tested in clinical trials for its ability to deliver drugs across the blood-brain barrier (BBB). This technique has the potential to treat neurological diseases by preferentially delivering drugs to targeted regions. Yet despite its potential, the physical mechanisms by which microbubbles alter the BBB permeability remain unclear, as direct observations of microbubbles oscillating in cerebral capillaries have never been previously recorded.

The purpose of this study was to reveal how microbubbles respond to ultrasound when within the microvessels of living brain tissue.

**Methods:** Microbubbles in acute brain slices acquired from juvenile rats perfused with a concentrated solution of SonoVue® and dye were exposed to ultrasound pulses typically used in BBB disruption (center frequency: 1 MHz, peak-negative pressure: 0.2–1 MPa, pulse length: up to 10 ms) and observed using high-speed microscopy at up to 10 million frames per second.

**Results:** We observed that microbubbles can exert mechanical stresses on a wide region of tissue beyond their initial location and immediate surroundings. A single microbubble can apply mechanical stress to parenchymal tissues several micrometers away from the vessel. Microbubbles can travel at high velocities within the microvessels, extending their influence across tens of micrometers during a single pulse. With longer pulses and higher pressures, microbubbles could penetrate the vessel wall and move through the parenchyma, refuting a previous assumption that microbubbles are confined to vessels. The probability of extravasation scales approximately with mechanical index, being rare at low pressures, but much more common at a mechanical index ≥ 0.6.

**Conclusions:** These observations provide important insight into microbubble dynamics in microvessels, and are critical leads into ultimately identifying the microbubble activities that lead to both safe drug delivery and into activities we seek to avoid, such as petechiae and other bioeffects.

## INTRODUCTION

On-going clinical trials are demonstrating that focused ultrasound and microbubbles can disrupt the blood-brain barrier (BBB) and deliver drugs in patients with Alzheimer’s disease, brain cancers, and other neurological conditions (1,2). In these patients, pre-formed lipid-shelled microbubbles with a diameter between 1 and 10 μm are systemically administered. Focused pulses of ultrasound are then applied to a target brain region, driving the microbubbles circulating within the vessels to expand and contract, somehow delivering drugs across the BBB (3).

Despite the promise of this method, the mechanisms through which oscillating microbubbles lead to drug delivery remain relatively poorly understood. Biological mechanisms have been investigated through post-mortem analysis and in vivo imaging of dye extravasation, which have shown drug delivery to be associated with disruption of tight junctions between endothelial cells (4,5), increased numbers of transcytotic vesicles (6,7), and vessel rupture (8,9).

However, these biological observations cannot link the drug delivery event to the microbubble behaviors which occur in the brain during ultrasound exposure (10). Direct observations of oscillating microbubbles within vessels have only been made within other tissue types, such as the chorioallontoic membrane of a chick embryo (11), cheek pouch of a hamster, and thin sections of the liver or muscle of rats (12). In rat ceca, vascular confinement produced significant damping of the oscillations, as well as asymmetric bubble behaviors and vessel wall motion (13). In the microvessels of rat mesentery, microbubble-induced vessel invagination and distension were observed, as well as microjetting (14).

Many of these studies used short (microsecond), relatively high-amplitude pulses (0.8 – 7.2 MPapk-neg at a center frequency of 1 MHz), with few consecutive frames captured per pulse. In contrast, drug delivery across the BBB is typically performed using lower amplitudes (<0.8 MPapk-neg) and much longer pulses (typically around 10 ms) (15–17), in which the bubbles likely behave very differently. While these studies provide important insight into bubble dynamics in tissues, their relevance to drug delivery across the BBB is therefore limited.

Acute brain slices are a common tool in neurophysiology research, enabling direct measurements of cellular activity within an intact live tissue environment. Juvenile slices are especially valuable due to their increased tissue viability in vitro and increased transparency (18,19).

Here, we use acute brain slices from juvenile rats perfused with microbubbles to present direct optical observations of microbubbles in brain microvessels at up to 10 million frames per second (fps), when exposed to therapeutically relevant ultrasound pulses.

## METHODS

### Slice preparation

Slices were prepared from Wistar wild-type juvenile rats (postnatal day 5-15, Charles River). Animals were culled via intraperitoneal injection of pentobarbital, followed by femoral exsanguination, in accordance with Schedule 1 of the Animals (Scientific Procedures) Act 1986. Shortly (<15 minutes) after cessation of circulation, the carcass was transcardially perfused with a concentrated solution of SonoVue (40 % v/v, Bracco), heparin (0.05 mg/mL), and either Evans Blue (4 mg/mL) or Blue India Ink (40 % v/v) in 0.9 % saline. A total fluid volume of around 3-5 mL was perfused.

The corpus callosum was transected and live cortical slices were obtained by sectioning the frontal lobes under ice-cold artificial cerebrospinal fluid (aCSF) using a vibratome (Campden Instruments 7000smz). Between sectioning and ultrasound experiments, slices were kept in aCSF at room temperature.

### Ultrasound

Brain slices were placed inside a Perspex box (depth: 2 cm, width: 5 cm) which had mylar walls on its front and back to let ultrasound propagate into and out of the box with minimal attenuation (Fig. 1). The top of the box was open, allowing objective lenses to be placed within it. A 5-mm-thick layer of 0.5 % agarose was placed at the bottom of the box, on which the brain slice was placed. Agarose allowed a free acoustic path, while remaining transparent enough to allow light from underneath. A 100-μm-thick glass cover slip was placed over each brain slice to keep it in place during sonication. box was filled with aCSF at room temperature and placed within a tank of deionized water. The apparatus was set on a vibration isolation table (Vision Isostation, Newport).

**Fig. 1.**
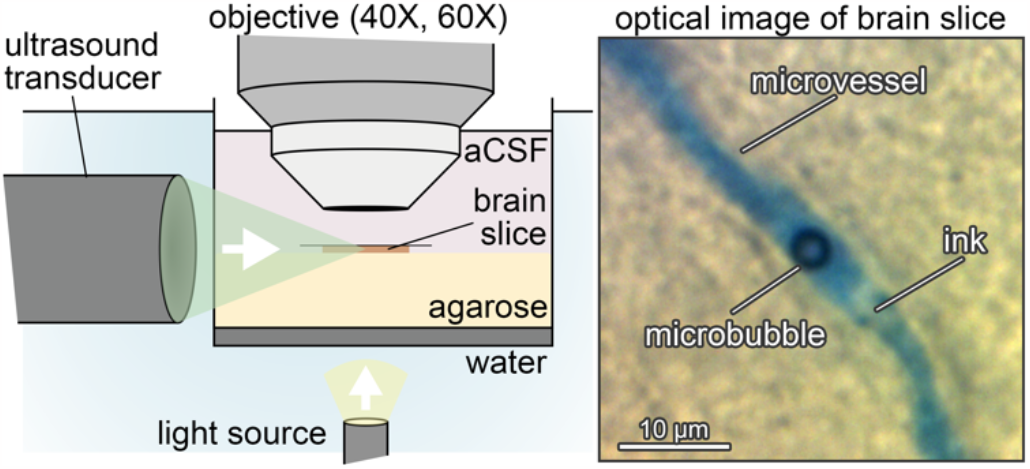
Experimental setup. (*Left*) A 250-μm-thick brain slice from a juvenile rat was immersed in artificial cerebrospinal fluid and placed between a light source and an objective (40X or 60X). A focused ultrasound transducer emitted sound onto the brain slice while videos were captured with either a color camera (1 frame per second (fps)), a high-speed camera (5,581 fps), or an ultra-high-speed camera (5 and 10 million fps). (*Right*) This experimental setup was able to capture images of microbubbles in brain microvessels.

Ultrasound pulses were emitted from an immersion transducer (A303 S-SU, diameter: 13 mm, focal distance: 15.2 mm, center frequency: 1 MHz; Olympus), driven by a function generator (33500 B Series, Agilent Technologies, Santa Clara, CA, USA) through a 50-dB amplifier (E&I) (Fig. S1). Ultrasound pulses were calibrated in situ before the start of each experiment using a needle hydrophone (diameter: 0.2 mm; Precision Acoustics), which was also used to align the focus of the transducer with the focal plane of the objective lens.

### Image acquisition

For the lower frame rate images (Fig. 2C, Fig. 4, Fig. 5ABEF), slices were imaged with a Chronos 1.4 camera (Kron Technologies Inc) connected to a 40x water dipping lens (LUMPLFLN, numerical aperture: 0.8, working distance: 3.3 mm; Olympus), illuminated from below with an LED light source (KL 2500, Schott) (Fig. S1 *left*). Videos were acquired at a frame rate of 5,581 fps, with a field of view of 608x400 pixels, and an exposure time of 174 μs. The pixel pitch was 0.16 μm. Images were simultaneously acquired using a color camera (ThorLabs DCC1645C CMOS, or IDS U3-3070CP-C-HQ; pixel pitch 0.08 μm) at 1 fps over a period of up to 2 minutes. This enabled clear images of the dye within the vessels before and after sonication to be obtained.

Ultra-high frame rate videos were acquired using an HPV-X camera (Shimadzu Corp) linked to a 60x water dipping lens (Fig. S1 *right*). The pixel pitch was 0.61 μm. Images were acquired at 10 Mfps, with ultrasound pulses of 15 cycles (Fig. 2AB), or at 5 Mfps with 60 cycle ultrasound pulses (Fig. 5C), although only approximately the first 50 were captured in the recording time in the latter instance. Illumination was provided by 10-ns synchronous laser pulses (CAVILUX Smart, Cavitar).

**Fig. 2.**
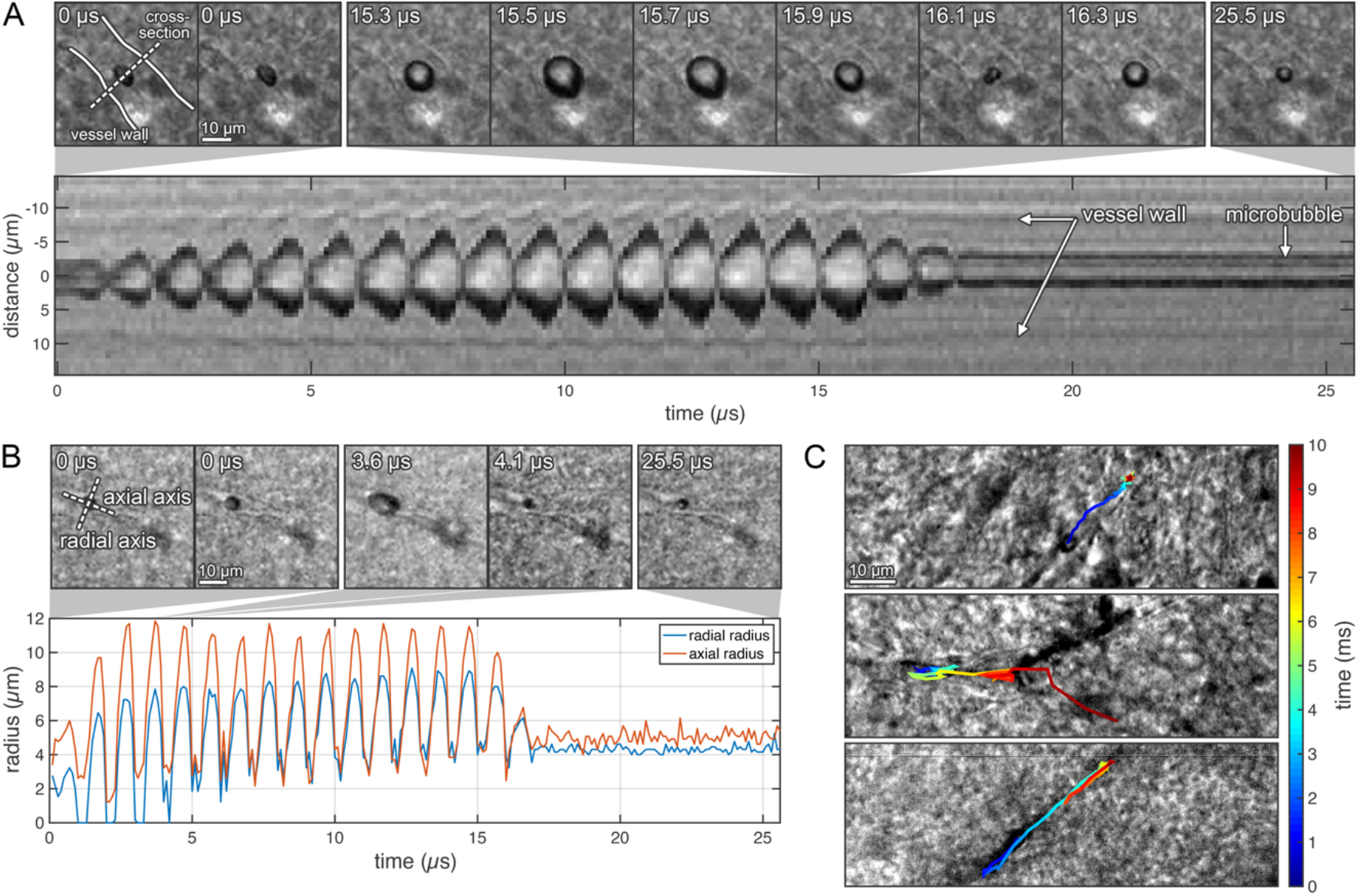
Microbubble dynamics during confinement in brain microvessels. (*A*) A microbubble’s oscillation can displace the vessel wall at the same rate. A microbubble in a microvessel was exposed to an ultrasonic pulse (center frequency (f_c_): 1 MHz, peak-rarefactional pressure (P_neg_): 0.8 MPa, pulse length (PL): 15 μs) and imaged at 10 million fps. (*Top*) The microbubble experienced a large expansion phase, followed by a sudden collapse. At 16.1 μs, an asymmetric collapse was observed in the shape of a ‘figure 8’. (*Bottom*) A streak of the microbubble’s cross-section (dotted white line in *Top*-*Left*) reveals a microbubble whose maximum expansion gradually increased with the number of exposure cycles. The vessel wall distended at the same rate as the microbubble, expanding with the bubble, but not invaginating as much when the bubble collapsed into itself. (*B*) A microbubble’s oscillations can be asymmetric. Two microbubbles were exposed to an ultrasonic pulse (f_c_: 1 MHz, P_neg_: 0.8 MPa, PL: 15 μs). The axial radius of one of the microbubbles was always larger than the radial radius. (*C*) A microbubble can move several microns within the microvessel. A microbubble was exposed to peak-rarefactional pressures of (*Top*) 0.6 MPa, (*Middle*) 0.4 MPa, and (*Bottom*) 0.6 MPa. The color represents the bubble’s location over the duration of the pulse. In all images, ultrasound propagated from left to right.

### Image processing

Videos were first motion-corrected in Matlab using a 2D cross-correlation algorithm to remove residual background motion caused by environmental vibrations or the direct effects of ultrasound on the tissue. The tissue deformation images in Fig. 4 were acquired by comparing selected frames before and during the pulse. A region of interest was chosen manually and compared using Digital Image Correlation software (Ncorr) (20). Bubble radii were measured by applying an intensity threshold to the images and fitting a circle using a Hough transform (21). This resulted in an uncertainty in the bubble diameter of approximately 0.2 μm, estimated from the variation in radius with chosen intensity threshold; this is close to the pixel pitch. The tracks of bubbles shown in Fig. 2C and 4B were acquired using the same algorithm, where the centroids of each circle in the image were also calculated.

### Image analysis

Measurements of vessel diameter (Fig. 2A) were made manually in Fiji by three independent observers (22). The errors calculated were the larger value out of the pixel pitch or the range of the independently measured values. Assessment of whether a bubble extravasated from a vessel was performed by two independent assessors who were blind to the ultrasound pressures applied, based on bubble motion, dye extravasation, and vessel shape changes.

## RESULTS

### Microsecond time-scale observations

Microbubbles in brain microvessels exposed to ultrasound expanded and contracted during the pulse (Fig. 2A, Video S1). These oscillations caused the vascular wall to move at the same rate as the bubble oscillation, at 1 million cycles per second. In the example shown in Fig. 2A, a microbubble with a diameter of approximately 5 μm in a microvessel with a diameter of 16.7 ± 0.6 μm was exposed to a 0.8-MPapk-neg pulse. The maximum and minimum vessel diameters after 10 cycles of oscillations were 19.4 ± 0.6 μm and 16.2 ± 0.6 μm respectively, showing that distension was greater than invagination. Interestingly, the bubble’s collapse did not result in collapse of the vessel, which contrasted with previous studies in larger vessels. The vessel diameter at the end of the recording, shortly after the end of the pulse, was 16.9 ± 0.9 μm. The uncertainty in these measurements was the larger value out of the pixel pitch or the standard deviation of the three values measured by three independent observers.

Several examples of microbubbles undergoing asymmetric oscillations were observed. In several instances, microbubbles expanded in an ellipsoidal shape, with more expansion along the axis of the vessel than radially. In one example, the bubble experienced a maximum to minimum diameter ratio of around 1.3 (parallel and perpendicular to the central axis of the vessel) (Fig. 2B, Video S2).

Microbubbles also experienced periodic hourglass-shaped oscillations, not associated with fragmentation. The axis of minimum contraction coincided with the central axis of the vessel lumen (Fig. 2A). These asymmetric shapes were not due to jetting, which can result in the penetration of one side of the bubble through the other (23). We have also confirmed experimentally that microbubbles can coalesce within small brain microvessels. As an example, two microbubbles were attracted to each other and subsequently coalesced over a short period of around 5 to 7 acoustic cycles when exposed to an 800-kPa ultrasound pulse (Fig. 3, Video S3).

**Fig. 3.**
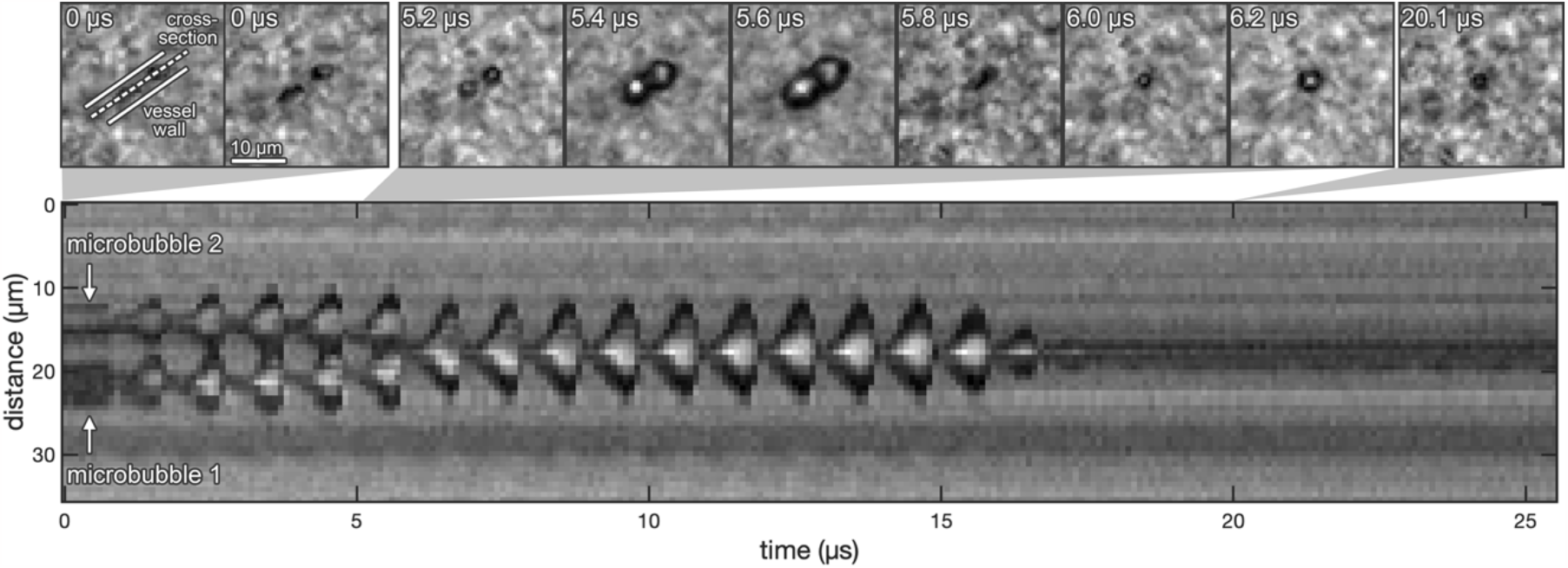
Microbubble coalescence in a brain microvessel. Two microbubbles in a microvessel were exposed to an ultrasonic pulse (f_c_: 1 MHz, P_neg_: 0.8 MPa, PL: 15 cycles) and imaged at ten million frames per second. (*Top*) Images of the microbubble were obtained before and during sonication. The bubbles were at first observed to be spatially separated, but had coalesced into a single bubble at a time point between 5.2 and 6.0 μs. (*Bottom*) A streak of the microbubble cross-sections (dotted white line in *Top-Left*) revealed microbubbles with spatially-separated oscillations that moved closer to each other. The two bubbles then merged and oscillated as a single bubble, but with larger-amplitude oscillations. A single bubble was observed after the ultrasound pulse.

### Millisecond-time scale observations

Microbubbles were also imaged at 5,581 fps to observe longer-timescale dynamics over the entire 10-ms pulse. During the 10-ms pulse, microbubbles could be driven tens of micrometers within the vessels (Fig. 2C). Bubbles typically moved along the vessels in the direction of ultrasound propagation, although this motion was often erratic (*Middle* of Fig. 2C, Video S4). Even when confined within vessels, microbubbles exposed to a 0.6 MPa_pk-neg_ pulse could achieve speeds of up to around 50 mm/s, as measured by the distance travelled between successive frames of the video.

In addition to the vessel wall oscillations observed at microsecond timescales, significant tissue displacements were also observed over millisecond timescales (Fig. 4, Video S5). Tissue displacements were quantified using digital image correlation (20), comparing the points in the tissue at their maximum displacement with an image from before the ultrasound pulse. Microbubbles could displace tissue located over 5 μm from the vessel lumen by up to around 0.5 μm.

**Fig. 4.**
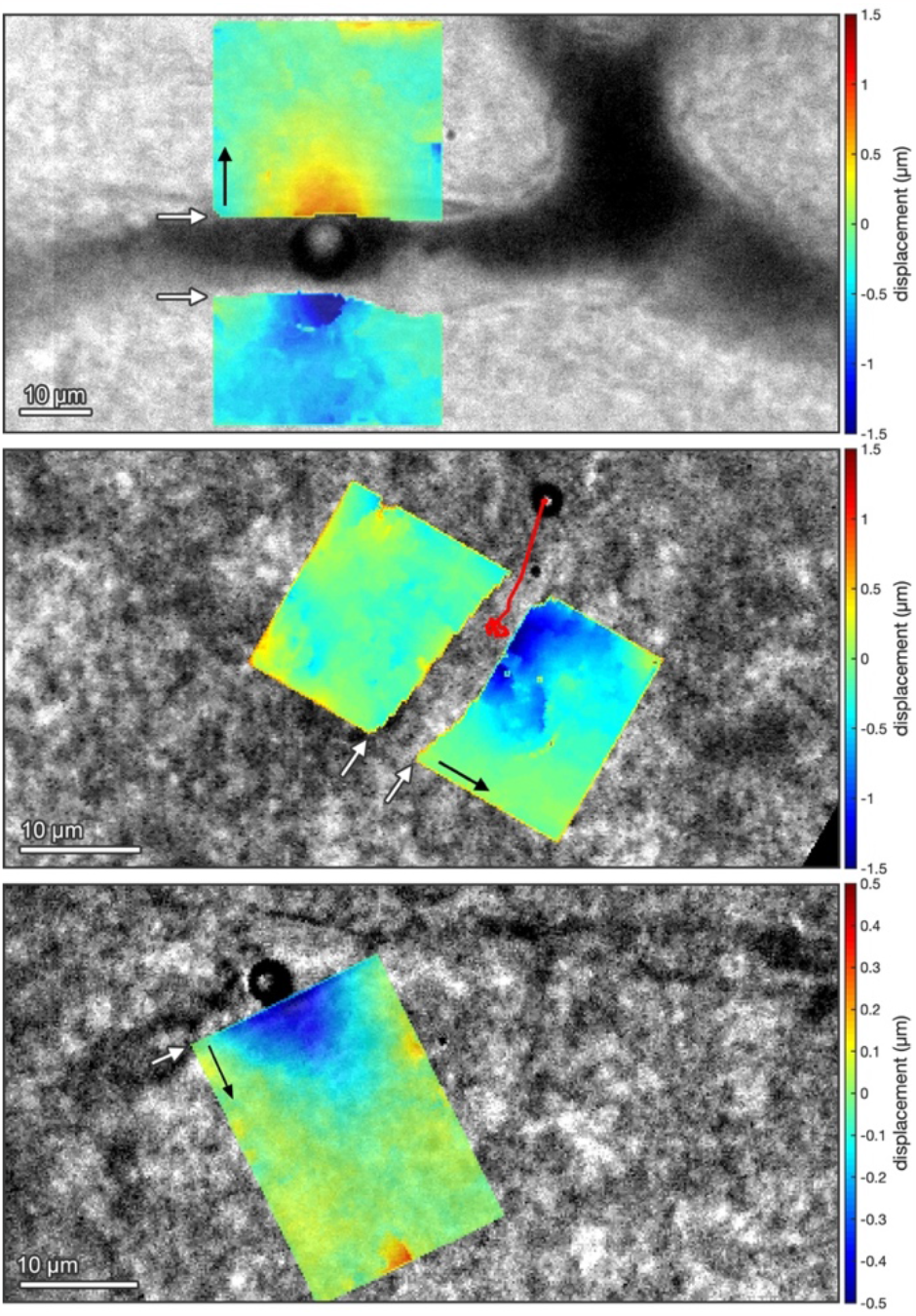
Tissue deformation in response to acoustic cavitation in brain microvessels. Microbubbles exposed to ultrasonic pulses (f_c_: 1 MHz, PL: 10 ms) produced tissue deformations well beyond the vessel wall on the longer, 10-ms timescale. The bubbles were exposed to (*Top*) 0.2, (*Middle*) 0.6, and (*Bottom)* 0.4 MPa_pk-neg_. The overlaid color maps show the maximum deformation of the tissue in micrometers compared to the initial location before the pulse. White arrows point to the vessel wall. Black arrows show the direction of positive displacement axis. Images were acquired at 5,581 frames per second. (*Middle*) The red line shows the motion of the microbubble during the pulse. In all images, ultrasound propagated from left to right.

While microbubbles have been shown to enhance extravasation of molecules from brain microvessels in vivo, we observed that, when exposed to ms-long pulses, microbubbles themselves can extravasate by passing through the microvessel walls. This can be associated with extravasation of the dye within the vessels (Fig. 5A), clearly indicating a puncture to the vessel. As shown in Fig. 5B (Video S6), after leaving the vessel, microbubbles can penetrate quite large distances (tens of micrometers) into the brain tissue.

**Fig. 5.**
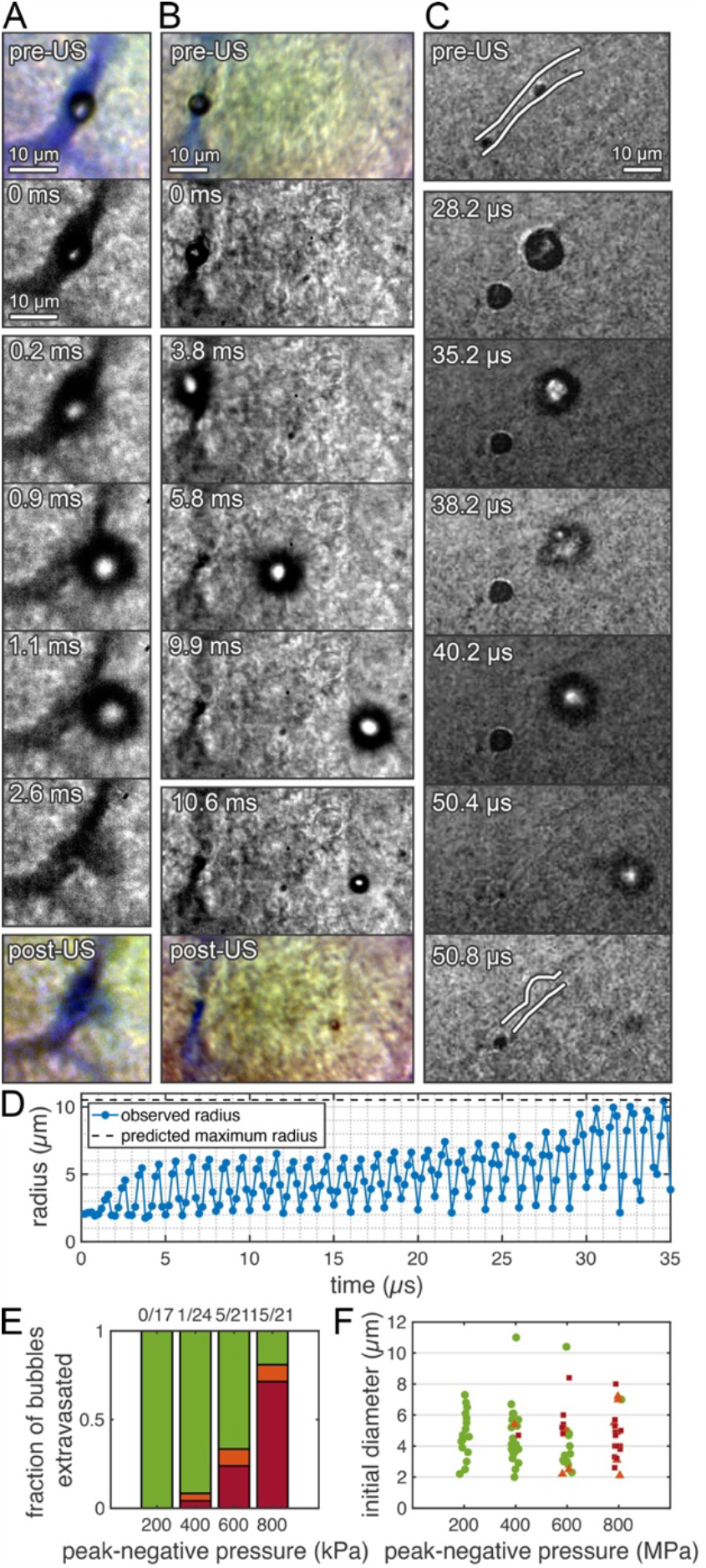
Extravasation of microbubbles due to the primary radiation force. Microbubbles in microvessels exposed to an ultrasonic pulse (f_c_: 1 MHz) extravasated in the direction of wave propagation. (*A*) A microbubble that extravasates can cause molecular delivery (video frame rate: 5,581 fps, P_neg_ 0.6 MPa, PL: 10 ms). The color images show the Evans Blue distribution before and after ultrasound exposure. (*B*) A microbubble can move within the vessel before extravasating and travel several tens of microns into the brain parenchyma (video frame rate: 5,581 fps, P_neg_ 0.4 MPa, PL: 10 ms). (*C*) A microbubble’s oscillation in confinement can evolve over time, while modifying the structure of the vessel itself, before extravasating (video frame rate: 5 Mfps, P_neg_ 1.0 MPa, PL: 50 μs). (*D*) Radius-time curve for the bubble in (*C*), showing low-amplitude oscillations while confined within the vessel, increasing gradually before it leaves the vessel. (*E*) Proportion of bubbles which extravasated (red), remained within the vessel (green), or whose behavior was unclear (orange) during exposure to a 10-ms pulse as a function of peak-rarefactional pressure. Extravasation did not occur at 0.2 MPa, was rare at 0.4 MPa, but becomes the dominant behavior at 0.8 MPa. (*F*) The initial radii of microbubbles counted may have influenced the likelihood of extravasation. In general, larger bubbles were more likely to extravasate at a given pressure, although this did not occur with very large bubbles (>10 μm diameter).

### Microbubble extravasation

Extravasation of microbubbles often imparted structural changes to the microvessels that persisted after the microbubble left the vessel (Fig. 5C, Video S7). Fig. 5D shows the radius-time curve of the bubble, which was exposed to a 1 MPa_pk-neg_ ultrasound pulse and imaged at 5 Mfps. Towards the end of the pulse, this bubble left the confines of the vessel. Over the first 15-20 cycles, the bubble initially oscillated with stable, moderate-amplitude oscillations (R_max_/R_0_ of around 3), before gradually increasing in amplitude (up to R_max_/R_0_ of around 5) as the vessel was deformed. The bubble then completely broke free of the confines of the vessel. At around the point at which the bubble left the vessel, its oscillations became asymmetric (Fig. 5C). The bubble fragments at this point, with the fragments continuing to move into the tissue.

We investigated the situations in which bubble extravasation occurs by observing the probability of extravasation at various ultrasound pressures. All microvessels from which bubbles extravasated were very small (around 5-10 μm in diameter). At mechanical indices (MI) that are generally considered safe and are close to the threshold for delivering drugs across the BBB (0.2-0.4), bubble extravasation was rare, with no instances observed at 0.2 and only one observed at 0.4 (Fig. 5E). However, the probability of observing this increased significantly with increasing pressure, with bubble extravasation being the dominant response observed in slices exposed to an MI of 0.8. At an MI of 0.4 and 0.6, extravasation tended to occur in larger bubbles (Fig. 5F). At 0.6, extravasating bubbles had diameters between 4.8 and 8.4 μm. At 0.4, the only bubble which extravasated was 6.1 μm in diameter. Very large bubbles, in excess of 10 μm, were rare, and did not extravasate.

## DISCUSSION

Drug delivery across the BBB using ultrasound and microbubbles has shown encouraging results in several clinical trials, yet the fundamental mechanisms behind the technique remain poorly understood due to a lack of direct observations of microbubbles within the microvessels of brain tissue. Here, we used acute brain slices from juvenile rats to directly observe microbubbles in microvessels when exposed to typical therapeutic ultrasound pulses, using high speed microscopy. We demonstrated that microbubbles can displace tissue beyond the wall of the vessel, extravasate from small microvessels, and oscillate asymmetrically under these conditions.

Microbubbles exposed to 10-ms ultrasound pulses can displace their surroundings on both microsecond and millisecond timescales. In contrast to previous studies in mesentery which demonstrated vessel distension was the dominant behavior on microsecond scales, we showed the majority of vessels exhibited more significant distension in response to microbubbles. This may be due to the smaller vessels, different tissue type, and longer pulse lengths used here (14).

At such high frequencies, the mechanical properties of vessels are unknown, but these results demonstrate that the vessel wall was sufficiently elastic to oscillate periodically over microsecond timescales. This force may stimulate mechanotransduction pathways, as has been shown directly in isolated cell cultures (24). Imaging at thousands of frames per second revealed displacement of tissues well beyond the vessel boundaries. Thus, while the focus on mechanisms have previously been on the vessel wall, one cannot ignore the mechanical stimuli that may be elicited in glial cells and neurons.

Microbubbles can extravasate from small microvessels under ultrasound parameters that have been used to deliver drugs across the BBB in animal and clinical studies (25,26). Microbubbles exited vessels and were pushed deep into the tissue by the primary radiation force. While vessel rupture has previously been described at much higher pressures than those used here (27,28), penetration deep into tissue has previously been observed in agarose gels, and only at higher PNPs in excess of 1 MPa_pk-neg_ at 1 MHz (29). This phenomenon is a plausible mechanism of drug delivery at higher ultrasound pressures. However, BBB opening has been demonstrated at pressures below the threshold at which we observed extravasation, indicating it is unlikely to be the sole mechanism (30).

Microbubble extravasation could be a mechanism of tissue damage, and, in particular, be a cause of localized red blood cell (RBC) extravasation. RBC extravasation has been observed in histological examinations of animals exposed to BBB opening treatments. The rate at which extravasation of microbubbles occurs in the slices correlates quite well with the relative rate of erythrocyte extravasation observed in *in vivo* studies of animals sonicated at these parameters, with a mechanical index of 0.2 and 0.4 being generally safe, but damage increasing at higher pressures (30–32).

Asymmetric oscillations of bubbles confined in small microvessels have been predicted theoretically, and our experimental observations support those predictions. This asymmetry in oscillations could alter the bubbles’ acoustic signatures, which may be significant in real-time therapy monitoring (33).

This study has several limitations when considering the relevance of its findings to a clinical or in vivo setting. Firstly, there was no blood flow or pressure in the vessels, key drivers of fluid extravasation, limiting our ability to observe dye leakage. The blood was replaced by SonoVue and dye; the presence of red blood cells in an in vivo setting may influence bubble dynamics (34). While we attempted to avoid bubble-bubble interactions, bubbles may have been present outside the focal plane.

Because the tissue was kept in vitro, cellular activity was likely lower than in an in vivo setting. However, the viability of acute slices is typically around 6-12 hours (18), considerably longer than the <2 hours these experiments took place over (see Supplementary Information).

Neonate animals were used due to the greater transparency and viability of their brain tissue over that of adults. While the tight junctions of the BBB are mostly developed in utero, many active transport mechanisms only develop in the weeks after birth (35). This may limit the potential of juvenile slices to fully investigate biological mechanisms of BBB disruption.

This study presents direct observations of microbubble dynamics in locally-intact brain tissue, when exposed to ultrasound pulses typical of those used to deliver drugs across the BBB. We observed asymmetric bubble oscillations, and tissue deformations that extend well beyond the endothelial wall. Bubbles are not stationary in an ultrasound field and can instead travel long distances due to the primary radiation force, applying mechanical forces to its surroundings as it moves. Single bubbles were also shown to puncture the vessel wall and apply mechanical forces throughout the brain parenchyma. These results enhance our understanding of the potential mechanisms of BBB disruption in vivo, which is critical to better interpreting the findings of ongoing clinical trials and to building a new generation of safer and more robust drug delivery technologies.

