## Supplementary Information for "Microbubble dynamics in brain microvessels"

### Supplementary Methods

#### Slice preparation

The aCSF solution contained 126 mM NaCl, 2.5 mM KCl, 1.25 mM $NaH_2PO_4$, 2 mM $CaCl_2$, 1 mM $MgCl_2$, 26 mM $NaHCO_3$, 15 mM glucose, and 5 mM pyruvate. Between sectioning and the ultrasound experiments, slices were kept in aCSF at room temperature.

It is typical in preparation of acute slices to saturate the extracellular solution with 95 % $O_2$:5 % $CO_2$. This reduces hypoxia in the tissue, as is especially important in adult or thick slices. In this study, the slices were instead maintained in aCSF in direct contact with atmospheric air. This was to avoid unwanted cavitation events unrelated to the SonoVue within the slices due to the gas saturation or trapping of small gas bubbles in or on the surface of the slice, which may affect the ultrasound experiments. Because juvenile slices are more resilient to hypoxia, and the slices were relatively thin (1), and because the experiments were completed in less than 2 hours post-mortem (limited by the lifetime of the microbubbles), the lack of $O_2$ saturation was unlikely to be a major concern.

#### Image processing

Videos were first motion-corrected in Matlab using a 2D cross-correlation algorithm to remove residual background motion caused by environmental vibrations or the direct effects of ultrasound on the tissue. To achieve sub-pixel motion correction, a small region was selected away from the bubble and spline-interpolated before comparing with the same region in the first frame to identify overall movement.

Bubble radii were measured by applying an intensity threshold to the images (a unique value was used for each image due to variations in overall image brightness) and fitting a circle to these binary images using a Hough transform in Matlab (3). This resulted in an estimated uncertainty in the bubble diameter of approximately 0.2 µm. This was estimated from the variation in radius with chosen intensity threshold, and is close to the pixel pitch. The tracks of bubbles shown in Fig. 2C and 5B were acquired using the same algorithm, where the centroids of each circle in the image were also calculated.

### Tissue Viability Assay

#### Methods

Dye was not observed outside vessels within tissue (except on the surface of the slice due to the sectioning process), indicating that the BBB was likely largely intact within the slices. In some videos of larger vessels, vasoconstriction was observed in response to the bubbles, again highlighting the continued activity of the cells in the vasculature.

Slice viability was assessed using triphenyltetrazolium chloride (TTC).(5) This is a marker for dehydrogenase activity associated with aerobic respiration. Viable tissue undergoing aerobic cellular respiration is turned deep red in the presence of TTC.

Slices were prepared in the same manner as those used in the main experiments, except that the ink in the perfusion was replaced with saline. This was to ensure the tissue was clear, enabling the red staining to be more easily observed and quantified.

Three tests were conducted. 5 slices were stained immediately after sectioning the brain. 5 slices were stained after immersion in 5 mL of aCSF for 2 hours at room temperature and open to the air. 5 slices were stained after immersion in PBS for 2 hours at room temperature while covered. The first two indicate the start and end of the typical recording period, while the latter was used as a negative control. For staining, 0.05 % of TTC was dissolved in PBS. The slices were placed in the TTC solution at 37 °C for 30 minutes.

The slices were imaged under a stereo microscope with an IDS color camera. The cortex of each slice was manually segmented in Fiji. The images were converted to HSV format in Matlab and the mean saturation value across all pixels of the cortex of each slice was used for comparison. Higher saturation values meant that the tissue appeared more intensely red.

**Results**

Acute brain slices are a well-established preparation to observe activity in living brain tissue.(6) However, the preparation methods here were unique to this set of experiments, and so a direct test of tissue viability on the samples used was performed using triphenyltetrazolium chloride (TTC).

The red intensity of the brain slices was significantly higher in the control and 2-hour aCSF groups than in the PBS group (Fig. S4). This indicated a good degree of viability of the slices at both the start and end of the recording period. However, there was quite significant variation in intensity, and small regions in most slices remained white. The results here focused on the mechanical properties of the tissues, meaning slightly reduced cellular activity was not a major concern (provided there was no significant necrosis). High viability may be essential for potential future experiments using this platform to investigate biological responses. This is therefore an area for future refinement.

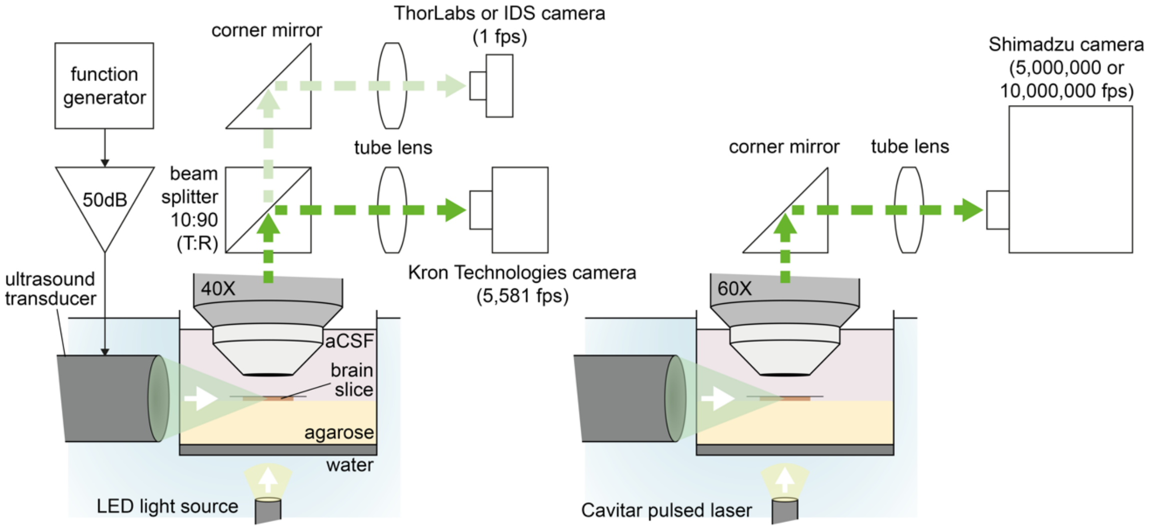


**Fig. S1. Experimental setups with optical and camera components.** A 250-µm-thick brain slice from a juvenile rat was immersed in artificial cerebrospinal fluid and placed between a light source and an objective. A focused ultrasound transducer emitted sound onto the brain slice, while videos were captured with one of two camera setups. (*Left*) In order to capture videos on the milliseconds timescale, light from an LED source was transmitted through a 40X objective and guided to a color camera at 1 frame per second (fps) and a high-speed camera at 5,581 fps. (*Right*) In order to capture videos on the microseconds time scale, light from a Cavitar 10-ns pulsed laser was transmitted through a 60X objective and guided into a Shimadzu camera at 5 or 10 million frames per second.

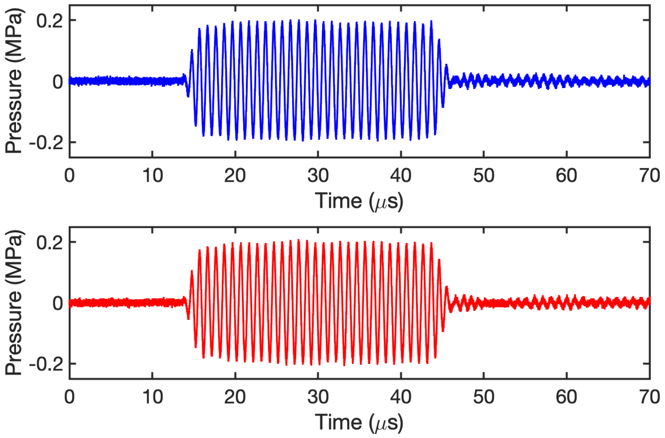


**Fig. S2. Shape of a 30-cycle ultrasound pulse.** Ultrasound pulses were measured using a needle hydrophone (*Top, Blue*) without a box present and (*Bottom, Red*) with the box present. The pressure amplitudes were very similar with or without the box. The ramp-up phase was short and only very low-amplitude ultrasound was present beyond the expected 30-cycle pulse waveform.

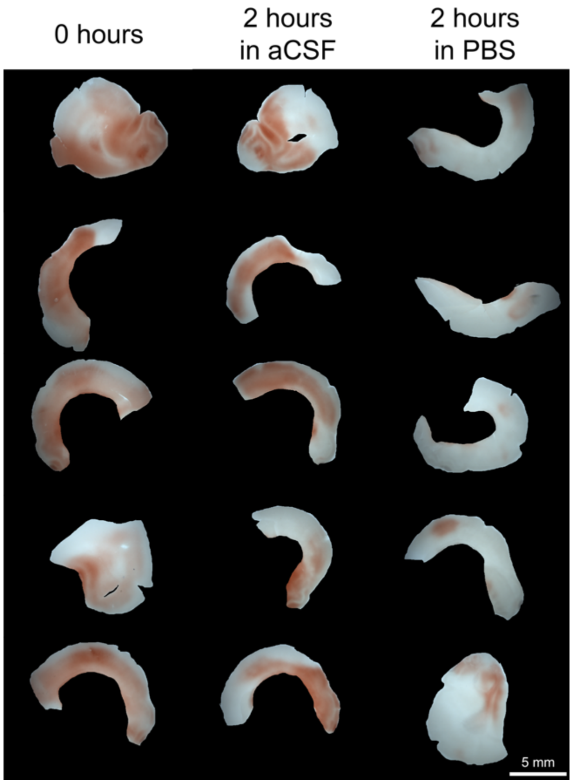


**Fig. S3. Triphenyltetrazolium chloride (TTC) staining of the brain slices.** The tissue is stained red when the tissue is undergoing aerobic respiration. The brain slices were stained with TTC (*Left*) at 0 hours, (*Middle*) 2 hours in aCSF, and (*Right*) 2 hours in PBS. The 0-hour time point refers to the time at which a typical experiment started, while the 2-hour time point refers to the time at which a typical experiment ended.

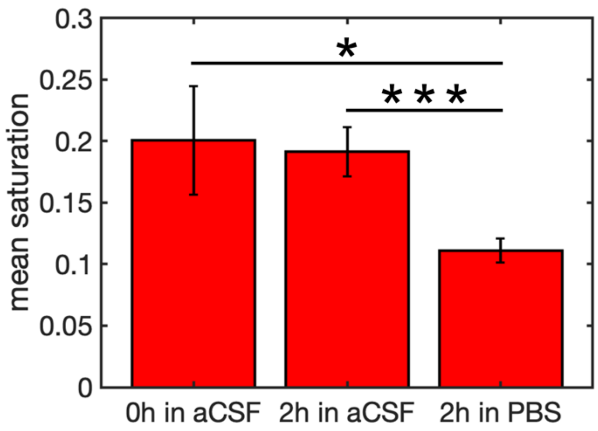


**Fig. S4. Mean TTC intensity for the cortex.** Error bars are standard deviations between slices. * p=0.04, *** p=8.4x$10^{-5}$

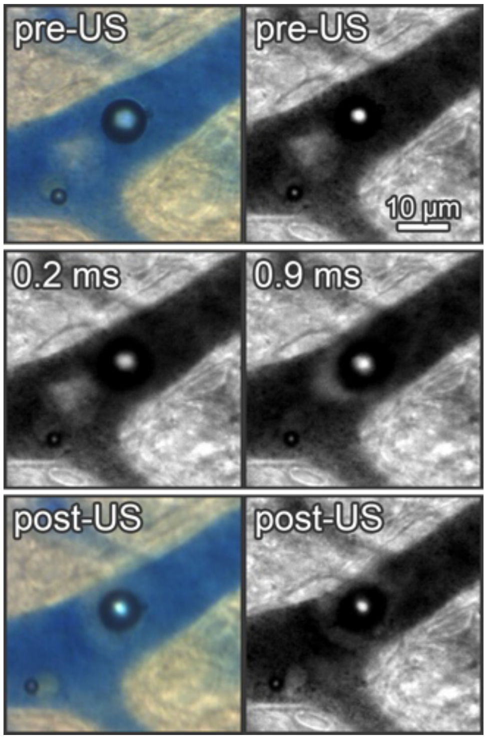


**Fig. S5. Microstreaming in a brain microvessel.** The brain slice was exposed to a 0.4-$MPa_{pk\text{-}neg}$, 10-ms pulse and imaged at 5,581 fps. Streaming patterns surrounding the microbubbles were observed during sonication. The region of dye around the bubble was affected by streaming, resulting in a clearly-defined boundary.

**Video S1. Acoustic cavitation in a brain microvessel.** A microbubble in a microvessel was exposed to an ultrasonic pulse (center frequency: 1 MHz, peak-rarefactional pressure: 0.8 $MPa_{pk\text{-}neg}$, pulse length: 15 cycles) and imaged at ten million frames per second. The vessel wall distended at the same rate as the microbubble, expanding with the bubble but not invaginating as much when the bubble collapsed into itself. At 16.1 µs, an asymmetric collapse was observed in the shape of a 'figure-eight'. Frames and a streak image of this video are presented in Fig. 2A.

**Video S2. Asymmetric oscillations in a brain microvessel.** Two microbubbles were exposed to an ultrasonic pulse (center frequency: 1 MHz, peak-rarefactional pressure: 0.8 $MPa_{pk\text{-}neg}$, pulse length: 15 cycles). The axial radius of one of the microbubbles was always larger than the radial radius. Frames of this video are presented in Fig. 2B.

**Video S3. Microbubble coalescence in a brain microvessel.** Two microbubbles in a microvessel were exposed to an ultrasonic pulse (center frequency: 1 MHz, peak-rarefactional pressure: 0.8 $MPa_{pk\text{-}neg}$, pulse length: 15 cycles) and imaged at ten million frames per second. The bubbles were first observed spatially separated, but then coalesced into a single bubble at a time point between 5.2 and 6.0 µs. Frames of this video are presented in Fig. 3.

**Video S4. Microbubble movement within a brain microvessel.** A microbubble in a microvessel was exposed to a 0.6-$MPa_{pk\text{-}neg}$, 10-ms pulse. The microbubble generally moved in the direction of wave propagation (left to right), but its motion was very erratic. A frame of this video and the bubble's travel path are presented in Fig. 2C.

**Video S5. A microbubble applying mechanical stress beyond the brain microvessel.** A microbubble exposed to ultrasonic pulses (center frequency: 1 MHz, pulse length: 10,000 cycles, PNP: 0.2 $MPa_{pk\text{-}neg}$) produced tissue deformations well beyond the vessel wall. The video was acquired at 5,581 frames per second. Ultrasound propagated from left to right. A frame and deformation maps for this video are presented in Fig. 4.

**Video S6. Microbubble extravasation on the millisecond time scale.** Extravasation of a microbubble during exposure to a 0.4-$MPa_{pk\text{-}neg}$, 10-ms pulse, imaged at 5,581 fps. Here, the bubble traveled up the vessel before exiting the vessel and traveling through the brain parenchyma. Frames of this video are presented in Fig. 5B.

**Video S7. Microbubble extravasation on the microsecond time scale.** Video showing extravasation of a microbubble exposed to a 1 $MPa_{pk\text{-}neg}$, 50-cycle pulse, imaged at 5 Mfps. Initially, the vessel became significantly distended and the bubble oscillation amplitude increased. The bubble then began transitioning out of the vessel and experienced highly asymmetric oscillations. The bubble then continued to penetrate into the tissue. There was a second bubble nearby which did not extravasate. Frames of this video are presented in Fig. 5C.

**Video S8. Microstreaming in a brain microvessel.** Microbubbles in a microvessel were exposed to an ultrasonic pulse (center frequency: 1 MHz, peak-rarefactional pressure: 0.4 $MPa_{pk\text{-}neg}$, pulse length: 10 ms) and imaged at 5,581 fps. Streaming was observed around both microbubbles in the vessel. The region of dye around the bubbles were affected by the streaming and had a clearly-defined boundary. Frames of this video are presented in Fig. S5.
